# XAB2 Dynamics during DNA Damage-Dependent Transcription Inhibition

**DOI:** 10.1101/2021.12.23.473962

**Authors:** Lise-Marie Donnio, Elena Cerutti, Charlène Magnani, Damien Neuillet, Pierre-Olivier Mari, Giuseppina Giglia-Mari

**Author notes:** To whom correspondence and request for materials should be addressed. G G-M. L.-M.D. and E.C. contributed equally to this work. **AUTHOR CONTRIBUTIONS**: P.-O.M. and G.G.-M. designed research; L.-M.D.; E.C.; D.N.; C.M. performed research; P.-O.M., E.C., L.-M.D. and G.G-M. analyzed data; and L.-M.D., E.C. and G.G.-M. wrote the paper.

## Abstract

Xeroderma Pigmentosum group A (XPA)-binding protein 2 (XAB2) is a multi-functional protein that plays a critical role in distinct cellular processes including transcription, splicing, DNA repair and mRNA export. In this study, we detailed XAB2 involvement during Nucleotide Excision Repair (NER), a repair pathway that guarantees genome integrity against UV light-induced DNA damage and that specifically removes transcription-blocking damage in a dedicated process known as Transcription-Coupled repair (TC-NER). Here, we demonstrated that XAB2 is involved specifically and exclusively in TC-NER reaction and solely for RNA Polymerase 2 transcribed genes. Surprisingly, contrary to all the other NER proteins studied so far, XAB2 does not accumulate on the local UV-C damage but on the contrary is remobilized after damage induction. This fast change in mobility is restored when DNA repair reactions are completed. By scrutinizing from which cellular complex/partner/structure XAB2 is released, we have identified that XAB2 is detached after DNA damage induction from the DNA:RNA hybrids, commonly known as R-loops, and from the CSA and XPG protein and this release is thought to contribute to the DNA damage recognition step during TC-NER. Importantly, we have disclosed a role for XAB2 in retaining RNAP2 on its substrate.

## INTRODUCTION

The DNA molecule, contained in the nucleus of our cells, forms the instruction manual for proper cellular functioning. Unfortunately, the integrity of our DNA is continuously challenged by a variety of endogenous and exogenous agents (e.g. ultraviolet light, cigarette smoke, environmental pollution, oxidative damage, etc …). These DNA lesions interfere with DNA replication, transcription and cell cycle progression and lead to mutation and cell death, which may cause cancer, inborn disease or aging (Chatterjee & Walker, 2017).

To prevent the deleterious consequences of persisting DNA lesions, all organisms are equipped with a network of efficient DNA repair systems. One of these systems is the Nucleotide Excision Repair (NER) which removes helix-distorting DNA adducts caused by ultraviolet light (UV) such as Cyclo-Pyrimidine Dimers (CPD) and 6-4 Photoproducts (6-4PP) (Giglia-Mari *et al*, 2011).

In mammals, the different steps of NER require about thirty different proteins that are recruited one by one into the DNA damage, as demonstrated by the different studies of NER proteins kinetics (Moné *et al*, 2004; van den Boom *et al*, 2004; Politi *et al*, 2005; Zotter *et al*, 2006; Rademakers *et al*, 2003). The first step of NER consists of damage recognition, followed by the opening of the DNA duplex, dual incisions on both sides of the damage, excision of 24–32 oligonucleotides containing the damage and, finally, gap filling by repair DNA synthesis.

NER is divided into two sub-pathways depending on where DNA lesions are located within the genome. The global genome repair (GG-NER) will detect and repair lesion throughout the genome, whereas transcription-coupled repair (TC-NER) is associated with RNA polymerase II (RNAP2) to repair lesions on the transcribed strand of active gene (Marteijn *et al*, 2014).

The NER system has been linked to rare human diseases classically grouped into three distinct NER-related syndromes. These include the highly cancer prone disorder Xeroderma Pigmentosum (XP) and the two progeroids diseases: Cockayne Syndrome (CS) and Trichothiodystrophy (TTD). Importantly, CS and TTD patients are not cancer-prone but present severe neurological and developmental features (Hanawalt, 1994).

Xeroderma Pigmentosum group A (XPA)-binding protein 2 (XAB2) is an evolutionarily highly conserved protein of 100 kDa and consists of fifteen tetratricopeptide repeat (TPR) motifs that play a role in protein-protein interactions. XAB2 protein was identified as a protein interacting with XPA, a NER factor, using a yeast two-hybrid system (Nakatsu *et al*, 2000). Next, it has been shown that this protein interacts also with the TC-NER-specific factors, CSA and CSB, as well as elongating RNAP2 (Nakatsu *et al*, 2000). XAB2 is also essential for early mouse embryogenesis as demonstrated by the preimplantation lethality observed in XAB2 knockout mice (Yonemasu *et al*, 2005).

Downregulation of XAB2 using either anti-XAB2 or siRNA specifically inhibited normal RNA synthesis and the recovery of RNA synthesis after UV irradiation (Nakatsu *et al*, 2000; Kuraoka *et al*, 2008). Furthermore, injection of anti-XAB2 in GG-NER deficient cells results in a significant reduction of UV-induced Unscheduled DNA Synthesis during repair (Nakatsu *et al*, 2000). These results suggest the involvement of XAB2 in transcription and in TC-NER.

Further studies have shown that XAB2 is a component of Prp19/XAB2 complex (Aquarius [AQR], XAB2, Prp19, CCDC16, hISY1 and PPIE) or Prp19/CDC5L-related complex required for pre-mRNA splicing (Kuraoka *et al*, 2008). XAB2, as well as PRP19 and AQR, has been involved in the DNA damage response (Onyango *et al*, 2016; Maréchal *et al*, 2014; Sakasai *et al*, 2017). Indeed, XAB2 is important for homologous recombination (HR) by promoting the end resection step (Onyango *et al*, 2016); PRP19 is a sensor of RPA-ssDNA after DNA damage (Maréchal *et al*, 2014) and AQR contributes to the maintenance of genomic stability via regulation of HR (Sakasai *et al*, 2017).

Moreover, AQR has also a role in the removal of R-loops, a three-stranded nucleic acid structure composed of a DNA:RNA hybrid and the associated non-template single-stranded DNA (Sollier *et al*, 2014). These structures can form during transcription when an RNA molecule emerging from the transcription machinery hybridizes with the DNA template. They are found abundantly in human gene promoters and terminators where RNA processing takes place (Wang *et al*, 2018).

Despite the knowledge acquired in the last decades on XAB2 and its different cellular roles, little is known on the exact crosstalk and dynamics between its diverse cellular functions, specifically between the DNA repair function, the involvement in transcription and its splicing activity. In this work, we described the molecular dynamics of XAB2 within the cell after UV-damage induction and during the TC-NER repair process. We determine *in vivo* that, in absence of XAB2, the transcription-coupled repair reaction is impaired and hence the transcription restart after UV damage is abolished. Surprisingly, unlike all the other NER proteins studied so far, the mobility of XAB2 is increased after DNA damage and XAB2 shows no accumulation on the local UV damage. This faster dynamic is not restored until DNA repair is completed and in damaged TC-NER-deficient cells, XAB2 remains more mobile. Interestingly, we demonstrate that, after DNA damage induction, XAB2 is not released from the splicing complex but is detached from R-loops, a recently identified substrate of XAB2 (Goulielmaki *et al*, 2021).

## RESULTS

### XAB2 is involved in TC-NER process

Two decades ago, Tanaka’s research group demonstrated the involvement of XAB2 in the NER pathway (Kuraoka *et al*, 2008; Nakatsu *et al*, 2000). However, the dynamics of XAB2 during the DNA Repair process remained to be elucidated. We aimed to study the molecular dynamics of XAB2 and the shuttling between its different functions when DNA damage is induced. Firstly, we wanted to verify that XAB2 is exclusively involved in the TC-NER reaction and not in other steps or pathways repairing UV-lesions.

The well-known standard assay used to quantify NER activity is the Unscheduled DNA Synthesis (UDS), which measure replication activity outside of the S-phase after UV treatment. This technique quantifies the refilling of the single-strand DNA gap by the DNA replicative machinery. When we performed UDS assay in XAB2 silenced cells, no decreased level of UDS was observed (as well as in mock-treated cells) (Figure 1A, blue and black columns and Figure S1). As a positive control, when we silenced the excision repair factor XPF, we observed a strong reduction in UDS level (Figure 1A, red column and Figure S1). This result shows that XAB2 is not involved in GG-NER sub-pathway, but does not exclude an involvement of XAB2 in TC-NER sub-pathway.

**Figure 1.**
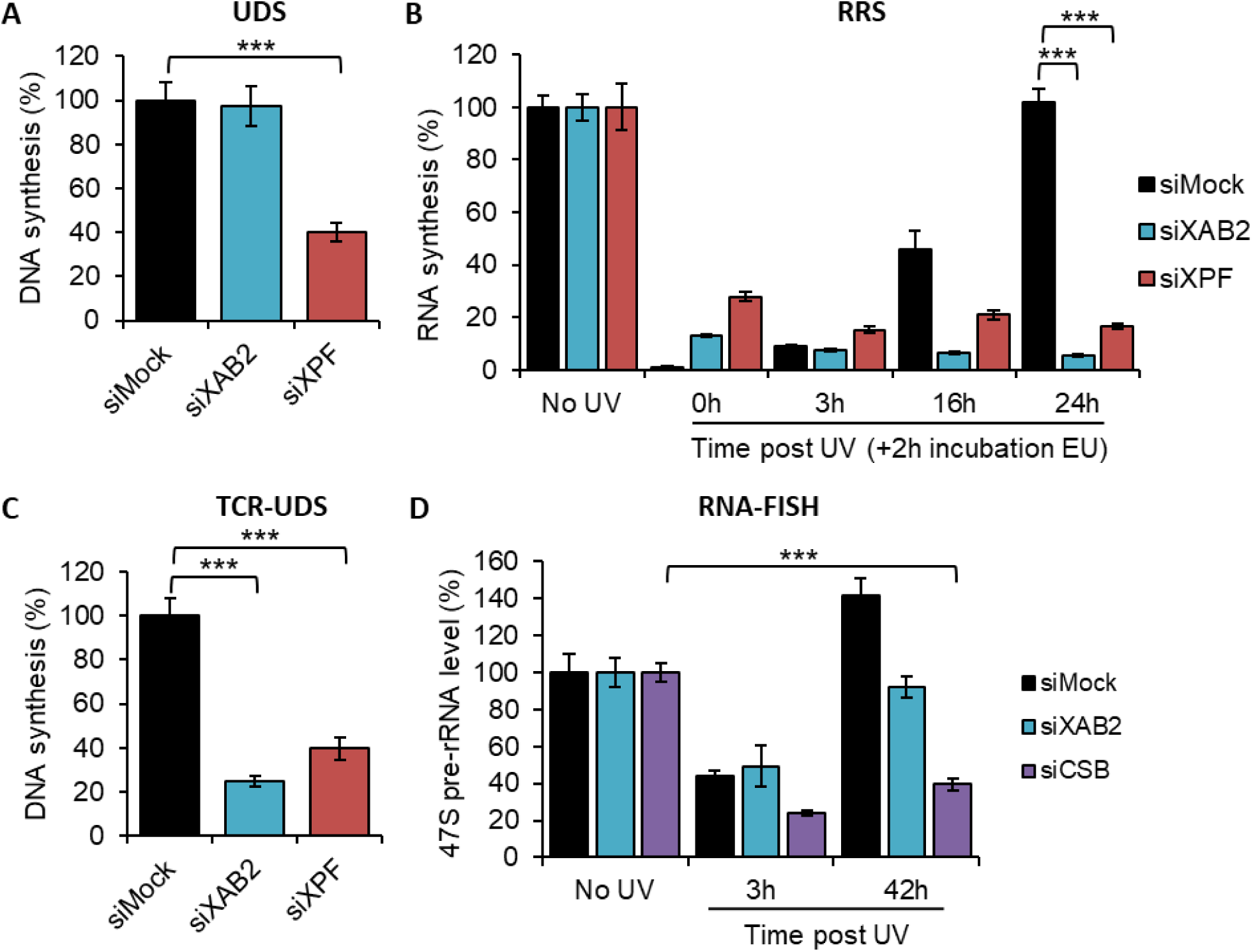
XAB2 involvement in DNA repair. **A)** Quantification of Unscheduled DNA Synthesis assay (UDS) determined by EdU incorporation after local damage (LD) induction with UV-C (100J/m^2^) in WT cells (MRC5 cells) treated with siRNAs against indicated factors. The graph is the average of two independent experiments and error bars represent the SEM obtained from at least 30 LDs. **B)** Quantification of RNA Recovery Synthesis (RRS) assay determined by EU incorporation after UV-C (10J/m^2^) exposure in WT cells treated with siRNAs against indicated factors. Error bars represent the SEM obtained from at least 50 cells and data are representative of three independent experiments. **C)** Quantification of TCR-UDS assay determined by EdU incorporation after LD induction with UV-C (100J/m^2^) in GG-NER deficient cells (XPC-/-cells) treated with siRNAs against indicated factors. Error bars represent the SEM obtained from at least 15 LDs and data are representative of two independent experiments. **D)** Quantification of RNA-FISH assay showing the 47S pre-rRNA level after UV-C (16J/m^2^) exposure in WT cells treated with siRNAs against indicated factors. Error bars represent the SEM obtained from at least 30 cells and data are representative of two independent experiments. For all graph, p-value of student’s test compared to No UV or siMock condition : ***<0.001.

The commonly used assay measuring TC-NER activity is the RNA Recovery Synthesis (RRS). In this assay, the newly transcribed RNA is measured via the incorporation of a nucleoside analog coupled to a fluorophore. The experiment is conducted at different time point after UV irradiation (0, 3, 16 and 24h), in order to quantify, 3h after UV damage, the decline in transcriptional activity and 16-24h after UV irradiation, the restart of transcriptional activity after DNA repair. When we performed this assay in XAB2 silenced cells no restart of transcription after UV damage was observed (Figure 1B, blue column and Figure S2A), as well as in siXPF-treated cells due to the inability to repair DNA lesions (Figure 1B, red column and Figure S2A) and in contrast with siMock-treated cells (Figure 1B, black columns and Figure S2A). Surprisingly, contrary to Tanaka’s group which observed a decreased of nascent RNA synthesis in absence of XAB2, we observed in our experiment an increase of transcription in siXAB2-treated cells without irradiation (Figure S2B) (Kuraoka *et al*, 2008; Nakatsu *et al*, 2000). This result demonstrated an involvement of XAB2 in the TC-NER sub pathway, but did not discriminate between a role in the repair reaction *per se* or in the restart of transcription after repair (RTR) (Mourgues *et al*, 2013).

In order to discriminate this point, we performed an assay designed previously in our group that specifically measures repair replication during TC-NER: the TCR-UDS assay (Mourgues *et al*, 2013). For this assay, we performed UDS assay in GG-NER deficient cells using XPC (Xeroderma Pigmentosum complementation group C) mutant cells (XP4PA-SV). The cells were transfected with siRNA and then locally irradiated with UV-C through a filter. In order to precisely localize DNA-damaged areas, a yH2AX co-immunofluorescence labeling was performed, and repair-replication was quantified. In siXPF-treated XPC-negative cells, both the GG-NER and the TC-NER pathways are compromised and as expected low TCR-UDS levels were observed compared to siMock-treated cells (Figure 1C, red and black column and Figure S3). Knockdown of XAB2 results in a decrease of TCR-UDS levels (Figure 1C, blue column and Figure S3). This result demonstrates a role of XAB2 in the repair reaction itself, its silencing preventing the DNA-synthesis associated with the excision of UV lesions on actively transcribed genes.

Recently we demonstrate that a fully functional NER mechanism is necessary for repair of ribosomal DNA (rDNA), genes transcribed by the RNA polymerase I (RNAP1) (Daniel *et al*, 2018). To investigate the involvement of XAB2 in the repair of ribosomal genes, the level of RNAP1 transcription was measured at different time points after UV irradiation by using a specific ribosomal RNA probe coupled to a fluorophore, as described previously (Daniel *et al*, 2018). This probe recognizes the 5’ end of the rDNA transcript, the 47S pre-rRNA (upstream from the first site cleaved rapidly during rRNA processing) (a sketch of the 47S is depicted in figure S4A). In siMock treated cells, we observed a decrease of 47S levels 3h after UV-C exposure and the restart of RNAP1 transcription 40h after irradiation (Figure 1D, black column and Figure S4C). The TC-NER-deficient siCSB-treated cells presented a low level of rRNA synthesis even 40h after UV-C exposure (Figure 1D, violet column and Figure S4B and S4C). A restart of RNAP1 transcription was observed in the absence of XAB2, as for the control (Figure 1D, blue column and Figure S4C).

All these results demonstrated a function of XAB2 in the TC-NER repair reaction specifically and exclusively for RNAP2 transcribed genes.

### XAB2-splicing complex mobility is released from the DNA damage area

XAB2 is included in a splicing complex composed of five other proteins, namely Aquarius (AQR), PRP19, CCDC16, PPIE and ISY1 (Kuraoka *et al*, 2008). In order to explore how the XAB2 splicing complex is behaving after local damage induction, the localization of XAB2, AQR, PRP19 and CCDC16 was revealed by immunofluorescence assays at different time points after local UV-irradiation of the cells. In this assay, the fluorescence signal from each protein in the damaged area (visualized by a co-staining of yH2AX) was compared to the rest of the nucleus. Unexpectedly, in contrast with all other NER proteins studied so far, we observed a relatively rapid (1 hour after UV-irradiation) release of XAB2, AQR, PRP19 and CCDC16 from the damaged area (Figure 2A and 2B). The localization of the XAB2-splicing complex is re-established after the completion of DNA repair reactions, when the transcription is fully restarted, 16h after irradiation (Figure 2B).

**Figure 2.**
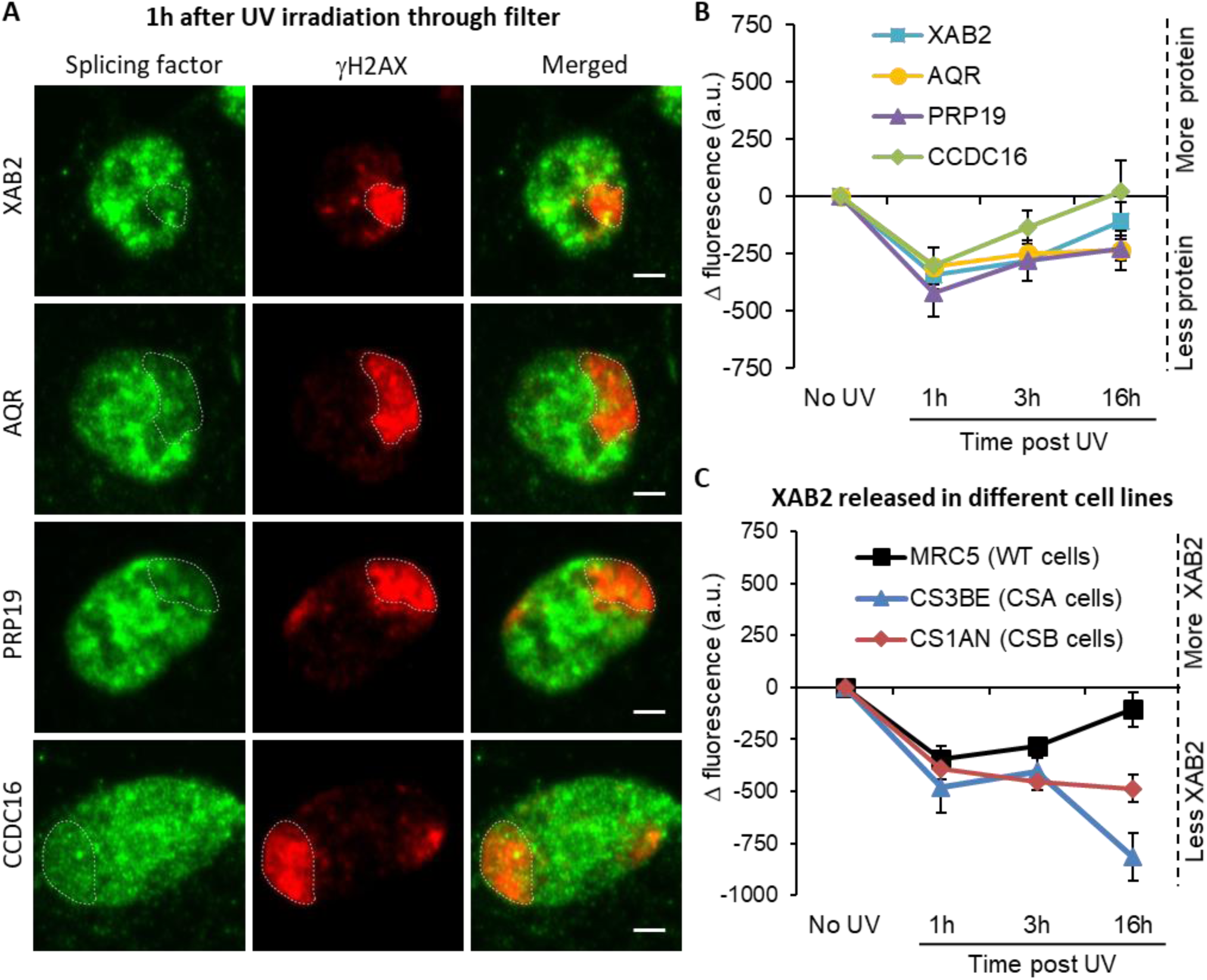
Splicing complex released from DNA damage. **A)** Representative confocal images of immunofluorescence against XAB2, AQR, PRP19 or CCDC16 (green) and γH2AX (red) one hour after local damage (LD) induction with UV-C (60J/m^2^). LD are indicated by dashed lines. Scale bar 3μm. **B)** Quantification of the immunofluorescence signal of the different splicing proteins on the LD after different times of recovery. **C)** Quantification of XAB2 signal on LD in different cell lines after different times of recovery. For both graph, the signal from the local damage has been subtracted from the background of each cell. Error bars represent the SEM obtained from at least 20 cells and data are representative of two independent experiments.

We thus verified if the entire XAB2-splicing complex was involved in TC-NER or whether only XAB2 played a role in this process. In order to measure the repair capacity of cells silenced for XAB2-related proteins, we performed UDS, TCR-UDS and RRS experiments in AQR/PRP19/CCDC16/PPIE/ISY1-siRNAs treated cells (Figure S5, S6, S7 and S8) and compared the results with XPF-siRNAs treated cell lines. Our results clearly show that none of the cells silenced for XAB2-related proteins are deficient in DNA Repair and both GG-NER (Figure S6) and TC-NER (Figure S7 and S8) are proficient in absence of AQR/PRP19/CCDC16/PPIE/ISY1.

In order to investigate whether XAB2 release from damaged areas was dependent of the TC-NER reaction, the localization of XAB2 was detected and quantified within locally damaged areas in TC-NER-deficient cells, CSA (CS3BE) and CSB (CS1AN) mutant cells. Interestingly, the absence of CSA and CSB did not hinder the release of XAB2 from locally damaged areas but this release continued 16h after UV-C exposure (Figure 2C blue and red curve compared to black curve and figure S9), suggesting that the re-establishment of the proper localization of XAB2 within the nucleus after the DNA repair process depends either on the repair process *per se* or on the restart of transcription after completion of the DNA repair reactions.

### XAB2 dynamic during TC-NER

To further analyze the mobility of XAB2 within the nuclei, we performed SPOT-FRAP (Fluorescent Recovery After Photo-Bleaching) experiments. In this technique, fluorescence molecules are photo-bleached in a small spot by a high intensity laser pulse; then the recovery of fluorescence within the bleached area is monitored over time (Figure S10A). With no treatment, this measure of fluorescence recovery corresponds to the protein mobility within the living cells (Figure S10A black curve). After perturbation of the nuclear environment (e.g. DNA damage), a protein can become less mobile if it physically interacts with a new substrate or a bigger complex (Figure S10A, green curve); more mobile if the protein is released from its substrate (Figure S10A, blue curve); or eventually will not change its mobility (Figure S10A red curve).

We stably transfected a vector expressing a fluorescently version of XAB2 (XAB2-GFP, Figure S10B) in different SV40-immortalized human fibroblast: wild-type cells (MRC5, Figure S10C), CSA deficient cells (CS3BE) and CSB deficient cells (CS1AN). In order to determine the minimum dose of UV-C needed to detect a significant difference in XAB2 mobility, MRC5 XAB2-GFP cell were irradiated with doses of UV-C ranging from 2 to 16 J/m^2^ and SPOT-FRAP experiments were performed at different time point after irradiation (Figure 3A). Interestingly, we observed a dose-dependent increase in mobility of XAB2 (Figure 3A). Doses of UV-C as weak as 2 and 4 J/m^2^ induced a moderate increase in mobility 3 hours post-irradiation and a re-establishment of the basal XAB2 mobility within 16 hours post-irradiation (Figure 3A, blue and yellow bar). High UV-C doses (16 J/m^2^) induced an increased mobility of XAB2 more rapidly (1-hour post-irradiation) and a re-establishment of the control mobility 24 hours post-irradiation (Figure 3A, red bar). At intermediate doses of 8 J/m^2^ of UV-C, we observed a significant increase in XAB2 mobility during repair (3h after UV-C exposure) and the following return to the normal condition once repair is completed and transcription restarted (16h after irradiation) (Figure 3A, green bar). Interestingly, in CSA and CSB mutant cells, the increase in XAB2 mobility is also observed after 8 J/m^2^ irradiation but lasted until 24h after UV-C exposure (Figure 3B) witnessing the fact that in these cells DNA damage is not repaired and hence initial XAB2 mobility is not restored. Interestingly, without damage, XAB2 mobility is reduced in TC-NER deficient cells compared to wild-type cells for still unknown reasons (Figure 3B, black histogram).

**Figure 3.**
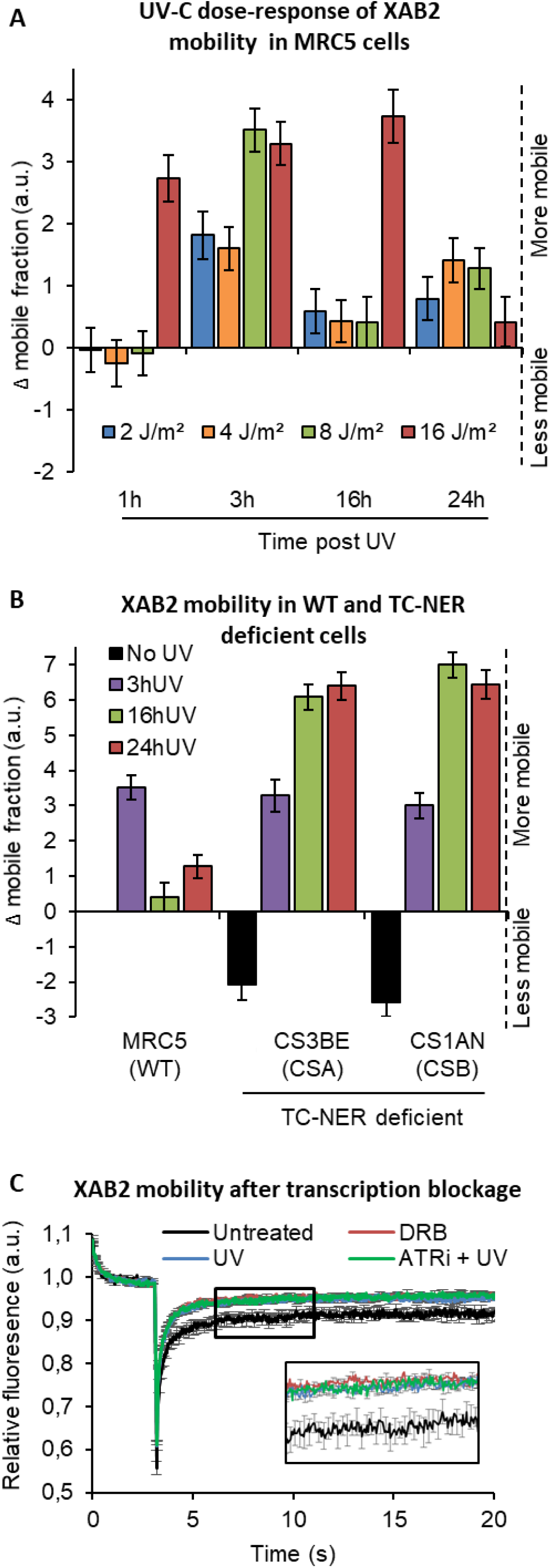
XAB2 dynamic during TC-NER. **A)** FRAP analysis of XAB2-GFP mobility in WT cells. Cells were treated or not with different doses of UV-C (2 to 16J/m^2^) and XAB2 mobility was measured at different time points after UV-C exposure. The No UV condition was used to calculate the change in bound fraction. **B)** FRAP analysis of XAB2-GFP expressed in WT cells (MRC5-SV) and TC-NER deficient cells (CSA -/- and CSB -/-). Cells were treated or not with 8J/m^2^ of UV-C. The No UV condition of the WT cell lines was used to calculate the change in bound fraction. **C)** FRAP analysis of XAB2-GFP mobility in WT cells. Cells were either untreated (dark line) or treated with 100μg/ml of DRB for 2h (red line) or with 10J/m^2^ of UV-C for 3h (blue line). Inhibitor of ATR pathway was added at 10μM in the medium 1h before irradiation (purple line). For all graph, error bars represent the SEM obtained from at least 10 cells and data are representative of two independent experiments.

The results of these experiments directed us to explore the possibility that the change in XAB2 mobility was due to transcription inhibition and not really to the repair process itself. In order to verify this hypothesis XAB2 mobility was measured after DRB (transcription inhibitor) treatment. The results show that XAB2 increased mobility in transcription inhibition conditions is very similar to the one measured upon UV treatment (Figure 3C).

As XAB2, the mobility of the late-stage spliceosomes change after UV irradiation and this mobilization depends on DNA damage response (DDR) signaling pathways (Tresini *et al*, 2015). Key mediators of DDR are the ATM and ATR kinases, which induced cell cycle arrest and facilitate DNA repair. To demonstrate that the change of XAB2 mobility is not due to the UV-Damage Response, we realized the same measures in presence of ATR and ATM inhibitors. Both drugs did not modify the increase of XAB2 mobility after UV irradiation (Figure 3C and Figure S10D).

Taken together, these results demonstrated that XAB2 mobility changes after DNA damage is triggered and sustained by the transcriptional inhibition.

### XAB2 is not released from the splicing complex during DNA repair reactions

The increase of XAB2 mobility after UV-induced transcription inhibition could be explained by either the release of XAB2 from a bigger complex and/or the release from an immobile (or nearly immobile) substrate such as the chromatin or a DNA-related substrate.

In order to distinguish between these two possibilities, we firstly investigated whether XAB2 dissociates after DNA damage induction from the splicing complex described earlier. XAB2 was immunoprecipitated together with AQR (Figure S11A). Interestingly, XAB2 was strongly and consistently immunoprecipitated 1-hour post-irradiation which corresponded to the time in which XAB2 mobility is increased and at the same time point more AQR is also immunoprecipitated. No clear release of XAB2 from AQR was observed at different time points.

In parallel, we also verified by PLA (Proximity Ligation Assay) whether the binding of XAB2 to AQR was modified after UV-C irradiation (Figure S11B). 2 hours after UV-C irradiation, instead of a release of XAB2 from AQR, we measured a stronger interaction (Figure S11B). However, this stronger interaction could very well result from the increase of AQR concentration 1h after UV irradiation (Figure S11C).

In conclusion, immunoprecipitation or PLA experiment could not explain the increased mobility of XAB2 after DNA damage, indeed XAB2 is not released from splicing complex during DNA repair reactions.

### XAB2 is released from R-loops during DNA repair reactions

Interestingly, while trying to immunoprecipitate XAB2 interacting partners during DNA Repair reactions, we could observe that systematically and consistently more XAB2 could be immuno-precipitated from nuclear extracts 1 to 3 hours after UV-C irradiation in WT cells (Figure S11A and 4A). At 16 hours post irradiation, we observed that the amount of XAB2 immunoprecipitated was comparable to the level observed in non-irradiated cells. Moreover, in CSA-/- and CSB-/-cell lines, in which UV-lesions on transcribed strands of genes are not repaired, the amount of XAB2 immunoprecipitated was remained high all along the time course of the experiment (3 and 16 hours) (Figure 4A). These results suggest that in TCR-deficient cells the binding between XAB2 and its substrate is not restored.

**Figure 4.**
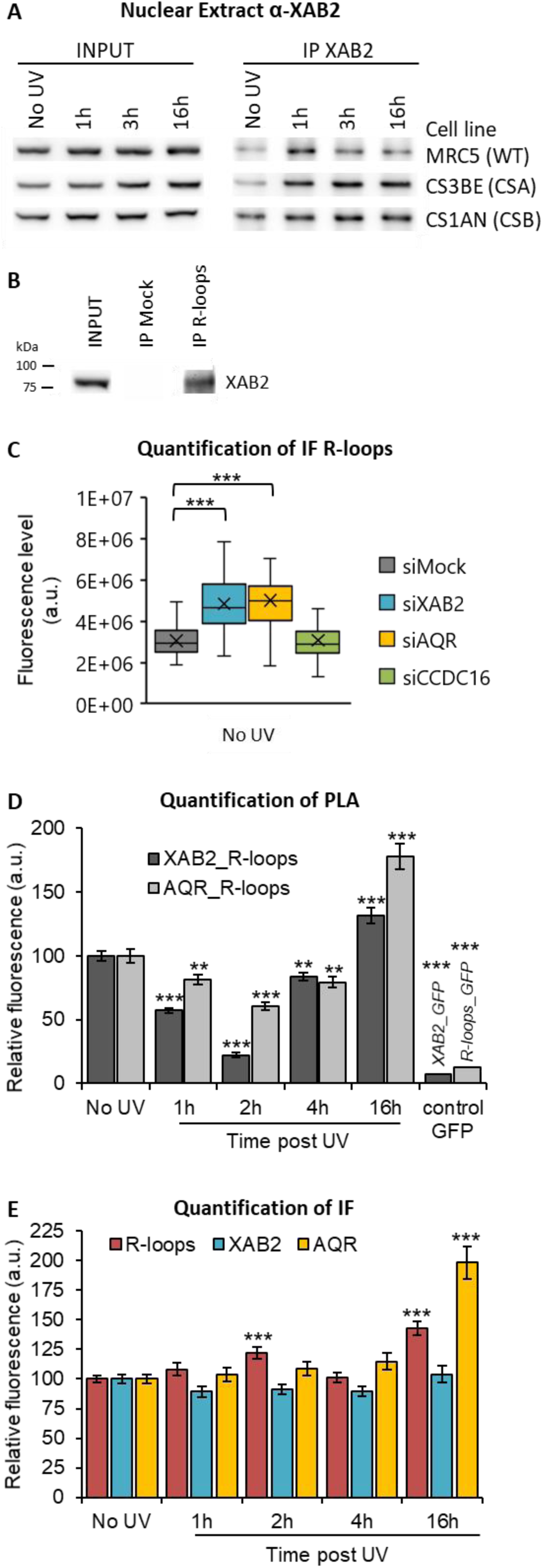
XAB2 and AQR are released from R-loop during DNA repair. **A)** Immunoprecipitation of XAB2 in nuclear extract from different cell lines treated with 10J/m^2^ of UV-C at different times. Bound proteins were revealed by Western blotting with antibodies against XAB2. INPUT, 10% of the lysate used for immunoprecipitation (IP) reaction. **B)** Immunoprecipitation of R-loops in non-crosslinked chromatin extract from WT cells. XAB2 bound to R-loops was revealed by Western blotting. INPUT, 10% of the lysate used for immunoprecipitation (IP) reaction. **C)** Quantification of immunofluorescence (from at least 50 cells) against R-loops in WT cells treated with siRNAs against indicated factors. **D-E)** Quantification of fluorescent signal in the nucleus against the couple XAB2_R-loops or AQR_R-loops from PLA experiment **(D)** or from the immunofluorescence done in parallel of PLA assay **(E)**. Error bars represent the SEM obtained from at least 50 cells. For all graph, data are representative of two independent experiments. For R-loops quantification, the nucleolar signal was subtracted from the nuclear signal. P-value of student’s test compared to No UV or siMock condition: **<0.01;***<0.001.

AQR has a role in the removal of DNA:RNA hybrids, commonly known as R-loops structure (Sollier *et al*, 2014) and recently XAB2 was found to be involved in the R-loops resolution (Goulielmaki *et al*, 2021). We thus decided to investigate if XAB2 has also a link with R-loops, a DNA-related substrate. Indeed, we could find by immunoprecipitation that XAB2 directly interacts with R-loops (Figure 4B) and that in cells silenced for XAB2 or AQR compared to control cells, we observed an accumulation of R-loops (Figure 4C and Figure S12A). We thus decided to examine whether XAB2 is released from R-loops after DNA damage induction by performing a PLA assay using a specific R-loops antibody (S9.6). We measured a strong and consistent reduction, more than 40% reduction, of the interactions between R-loops and XAB2 and between R-loops and AQR (Figure 4D and Figure S12B). This reduced interaction is not caused by a reduction in either R-loops, XAB2 or AQR concentration during DNA repair (Figure 4E and figure S12B). To verify that this result is specific for RNA-loops and not for a non-specific signal coming from the interaction of XAB2 with messenger RNAs, we performed the same assays (PLA and IF) in presence of RNAse H which specifically degrades R-loops structures and not the RNA (Figure S13A and S13B). Our results show that RNAse H decreases the quantity of RNA-loops (Figure S13C-F) and in doing so it decreases also the interaction with both XAB2 (Figure S13C and S13D) and AQR (Figure S13E and S13F). To verify whether XAB2 increase in mobility is due to a decreased interaction with the mRNA we performed PLA assays in UV-irradiated cells (Figure S14A and S14C). A decrease in the interaction between XAB2 and mRNA was observed (Figure S14A and S14C) but this decrease corresponded to the decrease of mRNA caused by UV-dependent transcription inhibition (Figure S14B and S14C) invalidating the results observed in the PLA XAB2-mRNAs (Figure S14A).

These results clearly demonstrate that XAB2 is released from R-loops during DNA repair reactions.

### XAB2 is released from CSA during DNA repair

Because XAB2 has been found to participate specifically in TCR-NER repair reactions (Figure 1C), we wanted to investigate whether part of the increased mobility of XAB2 observed after UV-induction was due to a release from repair complexes. We measured, by PLA, the amount of interactions between XAB2 and CSA, CSB, XPB and XPG proteins, observed during TC-NER. Amongst all the proteins tested, we could observe a clear and consistent release from the CSA protein 2h after UV irradiation (Figure 5A) and from the XPG protein 1h after UV irradiation (Figure 5C). The corresponding IF did not show a decreased quantity of CSA or XPG (Figure 5B and 5D) which validated the specificity of the XAB2-CSA and XAB2-XPG decreased interaction at those time points. On the contrary, no clear decrease in interaction was observed between XAB2 and CSB (Figure S15A and S15B) or XPB (figure S15C and S15D).

**Figure 5:**
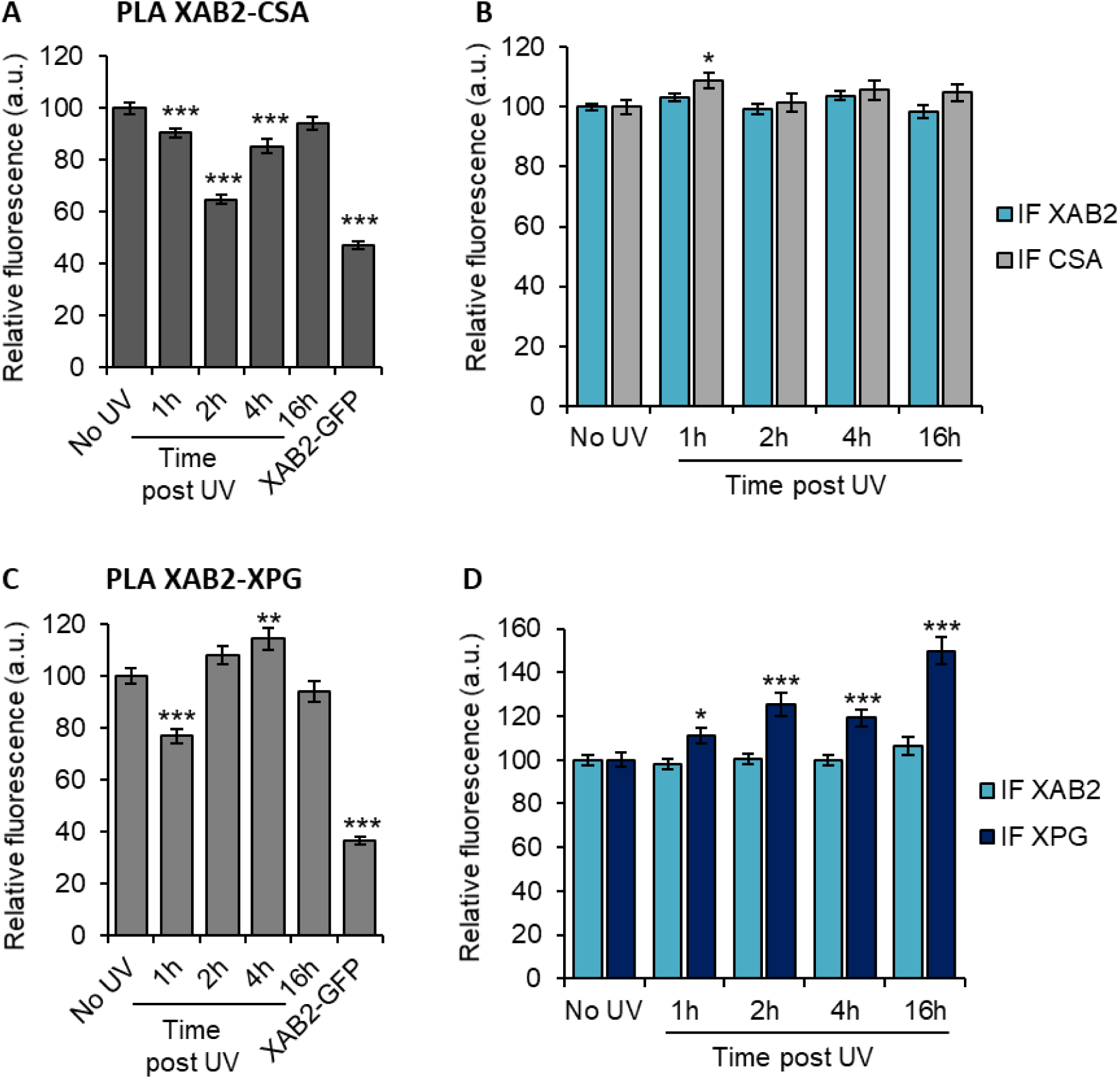
XAB2 are released from CSA and XPG during DNA repair. **A to D)** Quantification of fluorescent signal in the nucleus against the couple XAB2-CSA **(A and B)** and XAB2-XPG **(C and D)** from PLA experiment **(A and C)** or from the immunofluorescence done in parallel of PLA assay **(B and D)**. Data are the average of at least two independent experiments and error bars represent the SEM obtained from at least 80 cells. P-value of student’s test compared to No UV condition: *<0.05; **<0.01; ***<0.001.

### XAB2 depletion modify the mobility of the RNAP2

Because XAB2 was found to interact with RNAP2 (Nakatsu *et al*, 2000; Kuraoka *et al*, 2008) and because the increased mobility observed is dependent on the UV-dependent transcription inhibition step, we wanted to explore whether the depletion of XAB2 might play a role in the overall mobility of RNAP2. In order to investigate this point, we performed FRAP experiments on RNAP2-GFP expressing cells in presence and absence (depletion) of XAB2 (Figure 6) (Donnio *et al*, 2019). Interestingly, depletion of XAB2 greatly increase the mobility of RNAP2 (Figure 6, dark green curve versus dark blue curve). Moreover, UV-irradiation intensify this remobilization (Figure 6, light green curve versus light blue curve), demonstrating that XAB2 might play a role in maintaining the RNAP2 bound to the substrate in absence or presence of UV-lesions on the DNA.

**Figure 6:**
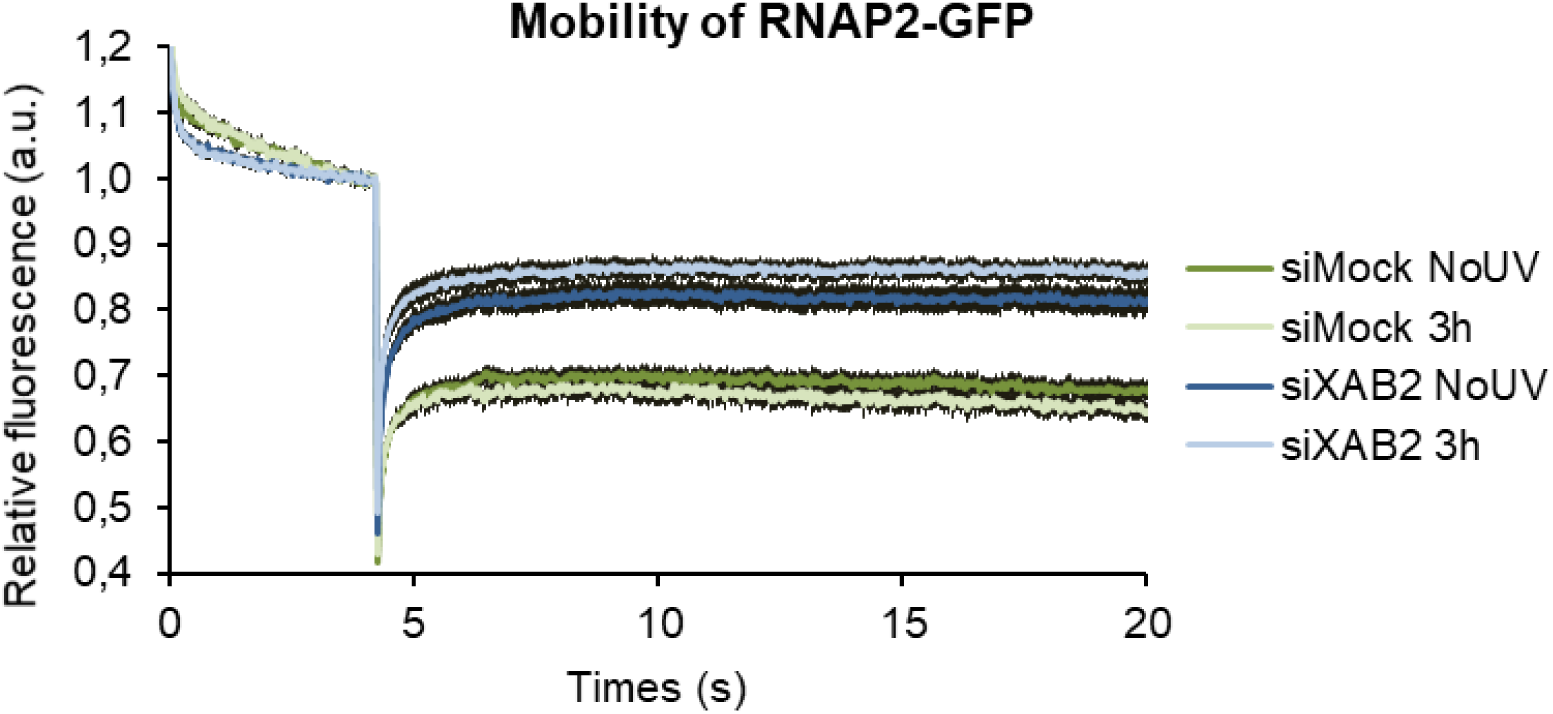
Pol2 behavior with XAB2. Strip-FRAP analysis of RNAP2-GFP expressing WT cells treated or not with UV-C (10J/m^2^) after siRNA mediated knockdown of the indicated factors. Error bars represent the SEM obtained from at least 10 cells and data are representative of two independent experiments.

## DISCUSSION

Helix distorting lesions continuously challenge the cell survival by interfering with and blocking fundamental cellular functions, such as transcription and replication. In order to prevent the deleterious effects of these events, cells have developed different mechanisms to restore an undamaged DNA molecule and allow the restart of cellular processes. The importance of rapidly reestablishing perturbed cellular functions is underlined by the presence of a repair mechanism directly coupled with transcription, like the Transcription-Coupled Nucleotide Excision Repair (TC-NER).

Two phases can be distinguished during TC-NER events: (i) the actual repair reaction of the damaged strand via the TC-NER sub-pathway and (ii) the resumption of transcription after repair. The experiment known as ‘RNA Recovery Synthesis’ (RRS) allow measuring the restart of transcription after DNA repair completion and thus the involvement of a protein in the TC-NER process. However, this assay does not discriminate between the two phases of the TC-NER. By consequence, proteins that are fully proficient in the repair reaction but fail in the transcription restart after DNA repair completion might have the same defect in RRS as proteins that are fully deficient in the DNA repair reaction.

Up to now, all studies have demonstrated the involvement of XAB2 in TC-NER process using only RRS experiments (Nakatsu *et al*, 2000; Kuraoka *et al*, 2008). In this study, we used a specific test called TCR-UDS, developed in house (Mourgues *et al*, 2013), and demonstrated that indeed XAB2 is solely needed for the repair reaction *per se* (Figure 1B and 1C).

XAB2 has been found as part of the pre-mRNA splicing complex composed of AQR, PRP19, CCDC16, PPIE and ISY1 (Kuraoka *et al*, 2008). However, none of these proteins seems to take part in the repair process, underlying the peculiar function of the splicing factor XAB2 in TC-NER repair (Figure S5).

We also demonstrated that, unlike all the other NER protein studied so far, XAB2 protein is released from the damage induced by UV-C exposure (Figure 3). Concomitantly, we also observed an increased XAB2 cellular mobile fraction after UV irradiation (Figure 4D). This release from transcription-blocking DNA lesions and increased mobility is surprising and atypical for a repair protein. However, this behavior has also been observed for late-stage spliceosomes (Tresini *et al*, 2015) and could be explained by the importance for the cell to rapidly provide access to the repair machinery.

Surprisingly, the increased mobility of XAB2 after UV irradiation occurs also in absence of CSA and CSB protein, in presence of transcription inhibitors and in presence of ATR and ATM inhibitors (Figure 2, 3 and Figure S6). These results strongly suggest that XAB2 remobilization is independent from the repair process but it is more a result of the damage-dependent transcription inhibition. Accordingly with this result, the return of XAB2 normal mobility is CSA and CSB dependent, suggesting that a proper repair and re-establishment of the transcription process is needed to restore the mobility of XAB2.

Previously, we reported that a complete TC-NER mechanism is required to repair the UV-lesions present on active ribosomal DNA, genes transcribed by RNA Polymerase 1 (RNAP1) (Daniel et al. 2018). Notably, both Cockayne syndrome proteins (CSA and CSB) are implicated in this specific repair reaction, as well as the UV-stimulated scaffold protein A (UVSSA), a protein required for the stabilization of CSB specifically after UV irradiation (Higa *et al*, 2016). Interestingly, XAB2 is involved only in repair of RNAP2-transcribed genes and not in repair of RNAP1-transcribed genes (Figure 1D), confirming a probable specific interaction with RNAP2 and reinforcing the idea that RNAP1 and RNAP2 repair processes are distinct although they share common proteins.

FRAP experiments in CSA and CSB mutant cells show a more immobile XAB2 fraction than in WT cells (Figure 3B) already without any damage induction. Tanaka’s group found that XAB2 interacts *in vitro* with CSA and CSB protein in the absence of DNA damage (Nakatsu *et al*, 2000). In addition, several studies demonstrated an involvement of CSB in transcription regulation (Boetefuer *et al*, 2018). As both CSB and XAB2 are necessary during the transcription process, it is therefore possible that the absence of CSB will modify the mobility of XAB2. However, it is not excluded that in CSA and CSB mutant cells, a low level of unrepaired oxidative damage (de Waard *et al*, 2004) might interfere with the proper XAB2 mobility, eventually modifying the amount of R-loops within these cells. Nevertheless, it is difficult to precisely estimate the exact number of R-loops between different cell types and this hypothesis is difficult to clearly assess.

The UV-induced remobilization of XAB2 is not explained by its release from the splicing complex, as demonstrated by co-immunoprecipitation and PLA experiment, but from both a chromatin-specific structures: R-loops and from repair proteins, CSA and XPG (Figure 4 and 5). An R-loop is a three-stranded nucleic acid structure, composed of a DNA:RNA hybrid and the associated non-template single-stranded DNA. This structure arises naturally in organisms from bacteria to humans, and have a multitude of roles in the cell (Belotserkovskii *et al*, 2018). In this work, we show that R-loops are a substrate for XAB2 and after DNA damage induction the interaction between XAB2 and R-loops are strongly reduced, this might explain the increase mobility of XAB2 during the TC-NER reaction. However, we also show that XAB2 interactions with CSA and with XPG are reduced after UV irradiation and during repair reactions (Figure 5). So probably, a combination of reduced interactions of XAB2 results in an increased cellular mobility.

A study demonstrates that absence of AQR induces R-loops formation which are actively processed into DNA double-strand breaks by XPF and XPG, the nucleotide excision repair endonucleases (Sollier *et al*, 2014). Without any damage, we also observe an increased level of cellular R-loops in absence of AQR, additionally we observed the same increase in cells silenced for XAB2 (Figure 4B), suggesting an involvement of these two proteins in R-loops resolution, as recently also demonstrated by Goulielmaki *et al*, 2021.

Transcription process and R-loops formation are finely interconnected. Indeed, R-loops formation can cause transcription blockage but transcription blockage due to DNA damage appears to result in R-loop formation (Steurer & Marteijn, 2017; Mullenders, 2015). Moreover, after UV irradiation, transcription activity declines and mRNAs levels are drastically reduced (Figure S9). PLA assays show that, after UV irradiation, XAB2 and total RNAs interaction is reduced, however, because the total amount of RNAs is diminished after transcription block, we assume that XAB2_RNAs interaction is lessened because of the intrinsic reduced amount of RNAs. Differently from total RNAs, in our study, we observe that R-loops do not decrease in number after UV-damage and transcription inhibition (Figure 4). Therefore, the interaction XAB2_R-loops and AQR_R-loops is specifically hindered after UV-damage (Figure 4).

Because XAB2 interacts *in vitro* with CSA and CSB protein (Nakatsu *et al*, 2000) and recently studies have shown an interaction with XPG (Goulielmaki *et al*, 2021), we decided to verify whether these interactions were modified during the DNA repair process and the concomitant transcription inhibition. Our results show a clear and consistent reduction of interaction between XAB2 and CSA (Figure 5A and 5B), and between XAB2 and XPG (Figure 5C and 5D) but not between XAB2 and CSB or XAB2 and XPB (Figure S10). A working hypothesis would then be that XAB2 needs to be released from CSA and XPG to allow a proper repair reaction.

The physical interaction between XAB2 and RNAP2 has already been established (Nakatsu *et al*, 2000; Kuraoka *et al*, 2008), however the exact relation between XAB2 and RNAP2 has not been disclosed. In this study, we have clearly demonstrated by FRAP experiments that the mobility of RNAP2 is severely affected in the absence of XAB2, namely, RNAP2 is released from its substrate and its mobility is strongly increased (Figure 6). When cells are subjected to UV-damage, a small RNAP2 immobile fraction can be observed, probably reflecting paused RNAP2 molecules on DNA damaged sites. However, in the absence of XAB2 and in presence of a DNA damage, RNAP2 mobility is even more increased than without damage, confirming that XAB2 might stabilize RNAP2 onto the transcribed strands of active genes in presence of damage. This result leads to the hypothesis that XAB2 might collaborate with RNAP2 in the detection of damage during TC-NER and hence be implicated in the first step of this process.

In conclusion, we described here an increased mobility of the protein XAB2 during the DNA damage dependent transcription inhibition. This increased mobility might partly be explained by the release of XAB2 from its substrate R-loops and its partner CSA and XPG. Importantly, we demonstrated that XAB2 plays a role of anchoring RNAP2 to its substrate with and without DNA damage. The absence of XAB2 might hinder the overall transcription activity of the cells as well as it severely affects the TC-NER capacity.

## MATERIAL AND METHODS

### Cell culture and treatments

The cells used in this study were: (i) wild-type SV40-immortalized human fibroblasts (MRC5); (ii) XPC– deficient SV40-immortalized human fibroblast (XP4PA, GG-NER deficient); (iii) CSA–deficient SV40-immortalized human fibroblast (CS3BE, TC-NER-deficient); (iv) CSB-deficient SV40-immortalized human fibroblast (CS1AN, TC-NER-deficient); (v) MRC5 stably expressing XAB2-GFP; (vi) CS3BE stably expressing XAB2-GFP (vii) CS1AN stably expressing XAB2-GFP and (vii) MRC5 stably expressing RNAP2-GFP. Immortalized human fibroblasts were cultured in DMEM (sigma) supplemented with 1% of penicillin and streptomycin (Gibco) and 10% fetal bovine serum (Corning) and incubated at 37°C with 5% CO_2_.

XAB2-GFP and RNAP2-GFP stably expressing cell lines were produced by transfecting XAB2-GFP or RNAP2-GFP using FuGENE® 6 Transfection Reagent (Promega) according to the manufacturer’ protocol. Selection was performed with G418 at 2 mg/ml.

DNA damage was inflicted by UV-C light (254 nm, 6-Watt lamp). Cells were globally irradiated with a UV-C dose of 2, 4, 8, 10 or 16 J/m^2^ or locally irradiated with a UV-C dose of 60 or 100 J/m^2^ through a filter with holes of 5 μm of diameter (Millipore). After irradiation, cells were incubated at 37°C with 5% CO_2_ for different periods of time. Inhibitor of ATR pathway (VE821) was added at 10μM in the medium 1h before irradiation.

### Transfection of small interfering RNAs (siRNAs)

The small interfering RNA (siRNAs) used in this study are : siMock, Horizon, D-001206-14 (10nM); siXAB2, Horizon, L-004914-01 (20nM); siXPF, Horizon, M-019946-00 (10nM); siCSB, Horizon, L-004888-00 (10nM). The final concentration used for each siRNA is indicated in parentheses. All siRNA from Horizon are a pool of four different siRNA.

Cells were seeded in six-well plates and allowed to attach for at least 24h. Coverslip were added inside the well if needed for experiment. 24h and 48h after seeding, cells were transfected with siRNA using Lipofectamine® RNAiMAX reagent (Invitrogen; 13778150) or GenJet (Tebu-Bio), according to the manufacturer’ protocol. Experiments were performed between 24h or 72h after the second transfection. Protein knockdown was confirmed for each experiment by western blot.

### Recovery of RNA synthesis (RRS) assay

MRC5 cells were grown on 18 mm coverslips. siRNA transfections were performed 24h and 48h before RRS assay. RNA detection was done using a Click-iT RNA Alexa Fluor Imaging kit (Invitrogen), according to the manufacturer’s instructions. Briefly, cells were UV-C irradiated (10 J/m^2^) and incubated for 0, 3, 16 and 24 h at 37°C. Then, cells were incubated for 2 hours with 5-ethynyl uridine. After fixation and permeabilization, cells were incubated for 30 min with the Click-iT reaction cocktail containing Alexa Fluor Azide 594. After washing, the coverslips were mounted with Vectashield (Vector). The average fluorescence intensity per nucleus was estimated after background subtraction using ImageJ and normalized to not treated cells.

### RNA Fluorescence In Situ Hybridization (RNA FISH)

Cells were grown on 18 mm coverslips, washed with warm PBS and fixed with 4% paraformaldehyde for 15 min at 37° C. After two washes with PBS, cells were permeabilized with PBS + 0.4 % Triton X-100 for 7 min at 4° C. Cells were washed rapidly with PBS before incubation (at least 30 min) with pre-hybridization buffer : 15% formamide in 2X SSPE pH8.0 (0.3M NaCl, 15.7mM NaH_2_PO_4_.H_2_O and 2.5mM EDTA). 35 ng of probe was diluted in 70 μl of hybridization mix (2X SSPE, 15% formamide, 10% dextran sulphate and 0.5 mg/ml tRNA). Hybridization of the probe was conducted overnight at 37° C in a humidified environment. Subsequently, cells were washed twice for 20 min with pre-hybridization buffer and once for 20 min with 1X SSPE. After several washing with PBS, the coverslips were mounted with Vectashield containing DAPI (Vector). The probe sequence (5’ to 3’) is: Cy5-AGACGAGAACGCCTGACACGCACGGCAC. Images of the cells were obtained using a Zeiss LSM 780 NLO confocal laser scanning microscope and the following objective: Plan-Apochromat 63X/1.4 oil DIC M27.

### Unscheduled DNA synthesis (UDS or TCR-UDS)

MRC5 or XP4PA cells, WT or GG-NER-deficient cells respectively, were grown on 18 mm coverslips. siRNA transfections were performed 24h and 48h before UDS assays. After local irradiation at 100 J/m^2^ with UV-C through a 5 μm pore polycarbonate membrane filter, cells were incubated for 3 or 8 hours (UDS and TCR-UDS respectively) with 5-ethynyl-2’-deoxyuridine (EdU), fixed and permeabilized with PBS and 0.5% triton X-100. Then, cells were blocked with PBS+ solution (PBS containing 0.15% glycine and 0.5% bovine serum albumin) for 30 min and subsequently incubated for 1h at room temperature with mouse monoclonal anti-yH2AX antibody (Ser139 [Upstate, clone JBW301]) 1:500 diluted in PBS+. After extensive washes with PBS containing 0.5% Triton X100, cells were incubated for 45min at room temperature with secondary antibodies conjugated with Alexa Fluor 594 fluorescent dyes (Molecular Probes, 1:400 dilution in PBS+). Next, cells were washed several times and then incubated for 30 min with the Click-iT reaction cocktail containing Alexa Fluor Azide 488. After washing, the coverslips were mounted with Vectashield containing DAPI (Vector). Images of the cells were obtained with the same microscopy system and constant acquisition parameters. Images were analyzed as follows using ImageJ and a circle of constant size for all images: (i) the background signal was estimated in the nucleus (avoiding the damage, nucleoli and other non-specific signal) and subtracted, (ii) the locally damaged area was defined by using the yH2AX staining, (iii) the average fluorescence correlated to the EdU incorporation was then measured and thus an estimate of DNA synthesis after repair was obtained.

### Immunofluorescence

Cells were plated on 12 or 18 mm coverslips in order to reach 70% confluence on the day of the staining. After two washes with PBS, cells were fixed with 2% paraformaldehyde for 15 min at 37° C. Cells were permeabilized by three short washes followed by two washes of 10min with PBS + 0.1 % Triton X-100. Blocking of non-specific signal was performed with PBS+ (PBS, 0.5 % BSA, 0.15 % glycine) for at least 30 min. Then, coverslips were incubated with primary antibody diluted in PBS+ for 2 h at RT or overnight at 4°C in a moist chamber. After several washes with PBS + 0.1 % Triton X-100 (three short washes and two of 10min) and a short wash with PBS+, cells were incubated for 1h at RT in a moist chamber with secondary antibody coupled to fluorochrome (Goat anti-mouse Alexa Fluor® 488 [A11001, Invitrogen] or 594 [A11005] and Goat anti-rabbit Alexa Fluor® 488 [A11008] or 594 [A11012], 1/400 dilution in PBS+). After the same washing procedure with PBS instead of PBS + 0.1 % Triton X-100, coverslips were finally mounted using Vectashield with DAPI (Vector Laboratories).

The variation of fluorescence in the locally irradiated zone has been calculated as for the UDS experiment.

### Proximity Ligation Assay (PLA)

PLA experiments were done using Duolink™ II secondary antibodies and detection kits (Sigma-Aldrich, #DUO92002, #DUO92004, and #DUO92008) according to the manufacturer’s instruction. The cells were fixed and permeabilized with the same procedure as immunofluorescence. After blocking 1h at 37°C with the Blocking Solution from the kit, incubation of the primary antibodies diluted in Antibody Diluent was performed at 4°C overnight. Duolink™ secondary antibodies were added and incubated for 1h at 37°C. If secondary antibodies were in close proximity (<40 nm), they were ligated together to make a closed circle by the Duolink™ ligation solution. The DNA is then amplified (rolling circle amplification) and detected by fluorescence 594 (red dot). Coverslips were mounted using Vectashield with DAPI (Vector Laboratories).

### Cytostripping

To remove the background generated by some antibodies or EdU incorporation, the cytoplasm of the cells was removed before fixation. After two washes with cold PBS, cells were incubated on ice 5 min with cold cytoskeleton buffer (10mM PIPES pH6,8; 100mM NaCl; 300mM Sucrose; 3mM MgCl2; 1mM EGTA; 0,5% Triton X100) followed by 5mn with cold cytostripping buffer (10mM Tris HCL pH7,4; 10mM NaCl; 3mM MgCl2; 1% Tween 40; 0,5% sodium deoxycholate). After 3 gently washes with cold PBS, cells were fixed.

### Images acquisition

For RRS, images of the cells were obtained using an Andor spinning disk : Olympus IX 83 inverted microscope, equipped with a Yokaga CSU-X1 Spinning disk Unit and BOREALIS technology for homogeneous illumination. The acquisition software is IQ3.

For RNAFish, UDS, TCR-UDS and IF of splicing complex after local damage, images of the cells were obtained using a Zeiss LSM 780 NLO confocal laser scanning microscope and the following objective: Plan-Apochromat 63X/1.4 oil DIC M27 or 40X/1.3 oil DIC. The acquisition software is ZEN.

PLA and some IF has been performed on a Zeiss Z1 imager right using a 40x/0.75 dry objective. The acquisition software is Metavue.

All images were analyzed with ImageJ software.

### Primary antibodies used for IF and PLA

Primary antibodies used for Immunofluorescence and PLA experiments were anti-XAB2 (mouse, sc-271037 [Santa Cruz Biotechnology], 1/1000 dilution and rabbit, A303-638A [Béthyl], 1/500 dilution), anti-AQR (IPB160 rabbit, A302-547A [Béthyl], 1/500 dilution), anti-CCDC16 (rabbit, HPA027211 [atlas antibodies], 1/250 dilution), anti-PRP19 (rabbit, ab27699 [abcam], 1/500 dilution), anti-DNA:RNA hybrid clone S9.6 (mouse, MABE1095 [Merck Millipore], 1/100 dilution and rabbit, Ab01137-23.0 [Absolute antibody], 1/100 dilution), anti-CSA (rabbit, GTX100145 [genetex], 1/400 dilution), anti-XPG (rabbit, sc84663 [santa-cruz], 1/1000 dilution), anti-Pol2tot (rabbit, A300-653A [Béthyl] 1/400 dilution, anti-Pol2Ser2P (rabbit, ab5095 [abcam] 1/100 dilution) and anti-Pol2Ser5P (rabbit, 13523S [cell signaling] 1/1000 dilution).

### Fluorescence Recovery after Photo-Bleaching (FRAP)

FRAP experiments were performed on a Zeiss LSM 780 NLO confocal laser scanning microscope (Zeiss), using a 40x/1.3 oil objective, under a controlled environment (37°C, 5% CO2). A narrow region of interest (ROI) centered across the nucleus of a living cell was monitored every 20 ms (1% laser intensity of the 488 nm line of a 25 mW Argon laser) until the fluorescence signal reached a steady state level (after ≈ 2 s). The same region was then photo-bleached for 20 ms at 100% laser intensity. Recovery of fluorescence in the bleached ROI was then monitored (1% laser intensity) every 20 ms for about 20 seconds. Analysis of raw data was performed with the ImageJ software. All FRAP data were normalized to the average pre-bleached fluorescence after background removal.

Analysis of SPOT FRAP data was performed as follows (Figure S6). The average fluorescence (over all cells) of the No UV condition was subtracted from the average fluorescence of the UV treated conditions. The obtained difference between the two FRAP curves was then summed point by point, starting from the bleach up to the following 100 measurements i.e. the area between the curve of interest and the untreated condition curve.

### Protein extraction

Cells cultured in 10-cm dishes were harvested by trypsinization and the pellet was washed once with PBS supplemented with the Protease Inhibitor Cocktail (PIC, Roche). The extraction of nuclear proteins has been performed using the CelLytic™ NuCLEAR™ Extraction kit (Sigma-Aldrich) complemented with PIC. Protein concentration was determined using the Bradford method. The samples were diluted with Laemmli buffer (10% glycerol, 5% β-mercaptoethanol, 3% sodium dodecyl sulfate, 100mM Tris-HCL [pH - .8], bromophenol blue), and heated 95°C before loading on an SDS-PAGE gel.

### Co-immunoprecipitation

For co-immunoprecipitation, 10 μl of protein G magnetic beads (Bio-adembead, Ademtech) were used per IP. 1μg of anti-XAB2 antibody (rabbit, A303-638A, Bethyl) were bound to the beads in PBS with BSA (3%) during 2h at 4°C with rotation. 100 μg of nuclear extracts were then incubated with beads-antibodies complex for 2h at 4°C with rotation. After two washes at 100 mM salt, two washes at 150mM and one wash at 100mM, beads were boiled in 2x Laemmli buffer and eluted samples loaded on a SDS PAGE gel.

### RNA/DNA Hybrid IP

Non-crosslinked MRC5 cells were lysed in 85 mM KCl, 5 mM PIPES (pH 8.0), and 0.5% NP-40 for 10 min on ice. Pelleted nuclei were resuspended in RSB buffer (10 mM Tris-HCl pH 7.5, 200 mM NaCl, 2.5 mM MgCl2) with 0.2% sodium deoxycholate, 0.1% SDS, 0.05% sodium lauroyl sarcosinate and 0.5% Triton X-100, and extracts were sonicated for 10 min (Diagenode Bioruptor, 60 cycles high power, 10sec ON and 20 sec OFF). Extracts were then diluted 1:4 in RSB with 0.5% Triton X-100 (RSB + T) and subjected to IP with the S9.6 antibody overnight at 4°C. RNaseA was added during IP at 0.1 ng RNaseA per microgram genomic DNA. Then protein G dynabeads (Invitrogen) washed with RSB + T was added and incubated for 3h. Beads were washed 4x with RSB + T; 2x with RSB; and eluted in 2x Laemmli buffer for 10 min at 95°C before loading on SDS-PAGE.

### Western-blot

Proteins were separated on a SDS-PAGE gel composed of bisacrylamide (37:5:1), blotted onto a polyvinylidene difluoride membrane (PVDF, 0.45μm Millipore) The membrane was blocked in PBS-T (PBS and 0.1 % Tween 20) with 5 % milk and incubated for 2 h at RT or overnight at 4° C with the following primary antibodies diluted in milk PBS-T : Rabbit anti-XAB2, A303-638A (Bethyl) 1/1000. Subsequently, membrane was washed repeatedly with PBS-T and incubated 1 h at RT with the following secondary antibody diluted 1/5000 in milk PBS-T: Goat anti-rabbit IgG HRP conjugate (170-6515; BioRad). After the same washing procedure, protein bands were visualized via chemiluminescence (ECL Enhanced Chemiluminescence; Pierce ECL Western Blotting Substrate) using the ChemiDoc MP system (BioRad).

### Statistical analysis

Error bars represent the Standard Error of the Mean (SEM) of the biological replicates. Microsoft Excel was used for statistical analysis and plotting of all the numerical data. Statistics were performed using a test of student to compare two different conditions (siMock vs siRNA X or No UV irradiation vs after irradiation) with the following parameters: two-tailed distribution and two-sample unequal variance (heteroscedastic).

## ACKNOWLEDGMENTS AND FUNDING SOURCES

This work has been funded by Agence Nationale de la Recherche (ANR-14-CE10-0009), Institut National du Cancer (PLBIO17-043 et PLBIO19-126), La Ligue contre le Cancer (218398) et Électricité de France (contrat 218398).

## Notes

**COMPETING INTEREST STATEMENT**: The authors declare no conflict of interest.

